# Epigenetic regulation of Wnt7b expression by the *cis*-acting long noncoding RNA lnc-Rewind in muscle stem cells

**DOI:** 10.1101/2020.01.03.894519

**Authors:** Andrea Cipriano, Martina Macino, Giulia Buonaiuto, Tiziana Santini, Beatrice Biferali, Giovanna Peruzzi, Alessio Colantoni, Chiara Mozzetta, Monica Ballarino

**Author notes:** Corresponding authors: Monica Ballarino; Dept. of Biol. and Biotechnology “Charles Darwin”, Sapienza University, P.le Aldo Moro 5, 00185 - Rome, Italy, Chiara Mozzetta; IBPM-CNR, P.le Aldo Moro 5, 00185 - Rome, Italy. Co-first authors (Andrea Cipriano, Martina Macino). Andrea Cipriano: Dept. of Obstetrics & Gynecology, Stanford University, 94306 Stanford, CA, USA.

## Abstract

Skeletal muscle possesses an outstanding capacity to regenerate upon injury due to the adult muscle stem cells (MuSCs) activity. This ability requires the proper balance between MuSCs expansion and differentiation which is critical for muscle homeostasis and contributes, if deregulated, to muscle diseases. Here, we functionally characterize a novel chromatin-associated lncRNA, lnc-Rewind, which is expressed in murine MuSCs and conserved in human. We find that, in mouse, lnc-Rewind acts as an epigenetic regulator of MuSCs proliferation and expansion by influencing the expression of skeletal muscle genes and several components of the WNT (Wingless-INT) signalling pathway. Among them, we identified the nearby *Wnt7b* gene as a direct lnc-Rewind target. We show that lnc-Rewind interacts with the G9a histone lysine methyltransferase and mediates the *in cis* repression of *Wnt7b* by H3K9me2 deposition. Overall, these findings provide novel insights into the epigenetic regulation of adult muscle stem cells fate by lncRNAs.

## INTRODUCTION

The transcriptional output of all organisms was recently found to be more complex than originally imagined, as the majority of the genomic content is pervasively transcribed into a diverse range of regulatory short and long non-protein coding RNAs (ncRNAs) (Abugessaisa et al., 2017; Carninci et al., 2005; Cipriano and Ballarino, 2018). Among them, long noncoding RNAs (lncRNAs) are operationally defined as transcripts longer than 200 nucleotides, which display little or no protein coding potential. Since the initial discovery, numerous studies have demonstrated their contribution to many biological processes, including pluripotency, cell differentiation and organisms development (Ballarino et al., 2016; Fatica and Bozzoni, 2014). LncRNAs were demonstrated to regulate gene expression at transcriptional, post-transcriptional or translational level. The function and the mechanisms of action are different and primarily depend on their (nuclear or cytoplasmic) subcellular localization (Carlevaro et al., 2019; Rinn and Chang, 2012; Ulitsky and Bartel, 2013). A large number of lncRNAs localize inside the nucleus, either enriched on the chromatin or restricted to specific nucleoplasmic foci (Engreitz et al., 2016; Sun et al., 2018). In this location, they have the capacity to control the expression of neighbouring (*in cis*) or distant (*in trans*) genes by regulating their chromatin environment and also acting as structural scaffolds of nuclear domains (Kopp and Mendell, 2018). Among the most important functions proposed for *cis*-acting lncRNAs, there is their ability to regulate transcription by recruiting repressing or activating epigenetic complexes to specific genomic loci (Huarte et al., 2010; Maamar et al., 2013; McHugh et al., 2015; Wang et al., 2011).

In muscle, high-throughput transcriptome sequencing (RNA-seq) and differential expression analyses have facilitated the discovery of several lncRNAs that are modulated during the different stages of skeletal myogenesis and dysregulated in muscle disorders (Ballarino et al., 2016; Lu et al., 2013; Zhao et al., 2019). Although the roles of these transcripts have been partially identified, we are still far from a complete undestanding of their mechanisms of action. For instance, the knowledge of the lncRNAs impact on adult muscle stem cells (MuSCs) biology is partial and only a few examples have been characterized as functionally important. In the cytoplasm, the lncRNA Lnc-mg has been shown to regulate MuSC differentiation by acting as a sponge for miR-125b (Zhu et al., 2017). In the nucleus, the lncRNA Dum was found to promote satellite cells differentiation by recruiting Dnmts to the developmental pluripotency-associated 2 (Dppa2) promoter, leading to CpG hypermethylation and silencing (Wang et al., 2015). In mouse, we have previously identified several lncRNAs specifically expressed during muscle *in vitro* differentiation and with either nuclear or cytoplasmic localization (Ballarino et al., 2015). Among them, we found lnc-Rewind (**Re**pressor of **w**nt **ind**uction), a chromatin-associated lncRNA conserved in human and expressed in proliferating C_2_C_12_ myoblasts. Here, we provide evidence on the role of lnc-Rewind in the epigenetic regulation of the WNT (Wingless-INT) signalling in muscle cells. The Wnt transduction cascade has been demonstrated to act as a conserved regulator of stem cell function *via* canonical (β-catenin) and non-canonical (planar cell polarity [PCP] and calcium) signalling and dysregulation of its activity has been reported in various developmental disorders and diseases (Nusse and Clevers, 2017). In the muscle stem cell niche, Wnt signaling is key in coordinating MuSCs transitions from quiescence, proliferation, commitment and differentiation (Brack et al., 2008; Eliazer et al., 2019; Lacour et al., 2017; Le Grand et al., 2009; Parisi et al., 2015; Rudolf et al., 2016). Because of this central role, the Wnt pathway is supervised by several mechanisms and different works have shown that lncRNAs can modulate it both at transcriptional and post-transcriptional levels (Zarkou et al., 2018). Here, we provide evidence that lnc-Rewind associates with the H3K9 methyltransferase G9a to regulate the deposition of H3K9me2 *in cis* on the nearby *Wnt7b* gene. Our data show that lnc-Rewind expression is necessary to maintain *Wnt7b* repressed and to allow MuSCs expansion and proper differentiation.

## RESULTS

### Lnc-Rewind is a conserved chromatin-associated lncRNA expressed in satellite cells

In an attempt to uncover novel regulators of MuSCs activity, we decided to take advantage of the atlas of newly discovered lncRNAs, that we previously identified as expressed in proliferating muscle cells (Ballarino et al. 2015). Among them, we focused on lnc-Rewind, which is a lncRNA enriched in proliferating myoblasts and overlapping pre-miR-let7c-2 (mmu-let-7c-2) and pre-miR-let7b (mmu-let-7b) genomic loci (Figure 1A). An evolutionary conservation analysis performed by examining FANTOM5 datasets (Noguchi et al., 2017) revealed the existence of a conserved transcriptional start site localized in the syntenic locus (Figure 1B). RNA-seq (Legnini et al., 2017) (Figure 1B) and semiquantitative (sq)RT-PCR analyses (Figure S1A) confirmed that also in proliferating human myoblasts this region is actively transcribed to produce a transcript (hs_lnc-Rewind) which displays an overall ∼46% of (exonic and intronic) sequence identity (Figure S1B). This is a relatively high value of conservation for lncRNAs (Pang et al., 2006) and adds functional importance to the lnc-Rewind genomic locus.

**Figure 1:**
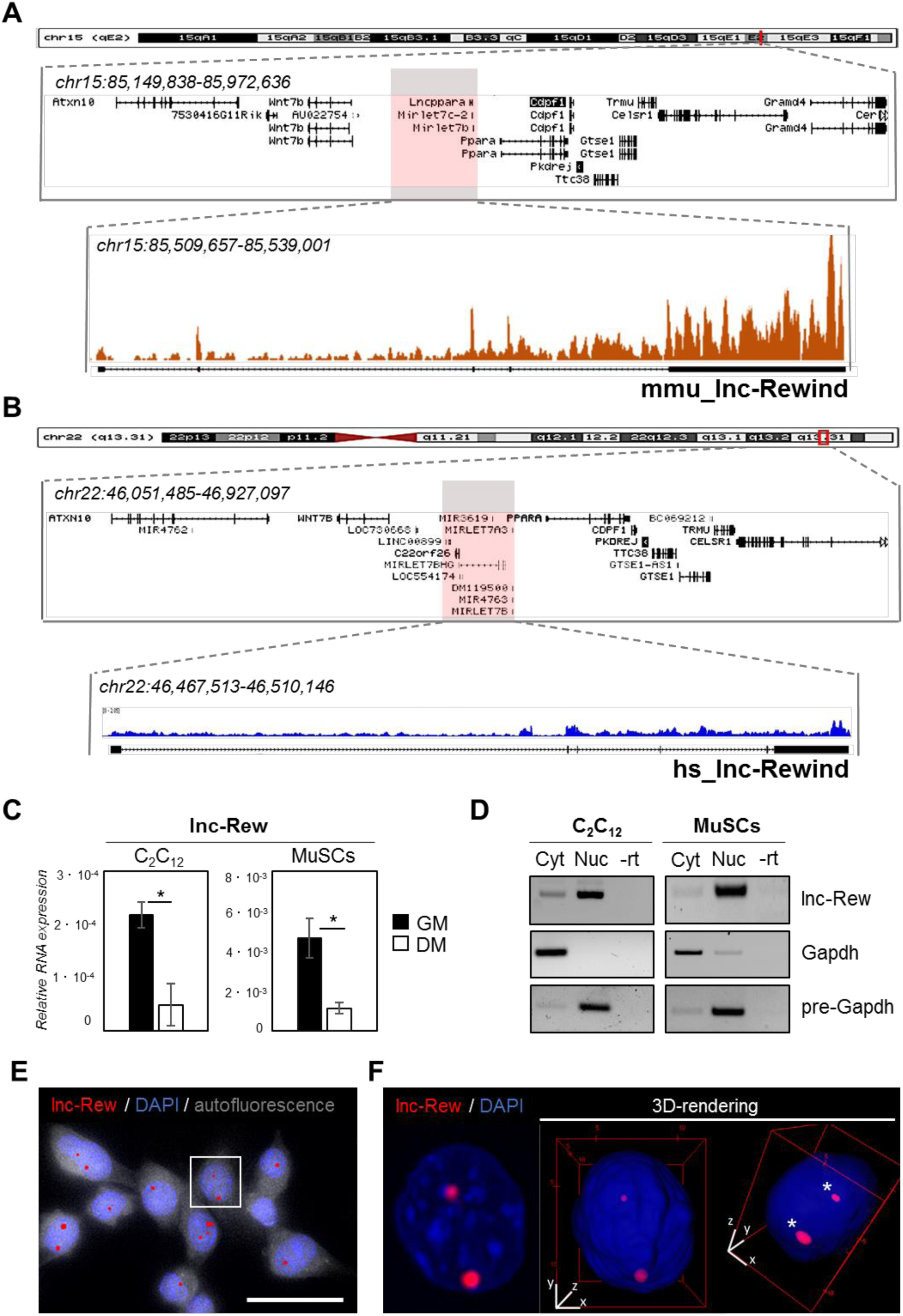
Lnc-Rewind is a conserved chromatin-associated lncRNA expressed in satellite cells. **A)** UCSC visualization showing the chromosome position and the genomic coordinates of lnc-Rewind (red box) in the mm9 mouse genome. The reads coverage from a RNA-Seq experiment performed in proliferating C_2_C_12_ cells (GM) (Ballarino et al. 2015; GSE94498) together with the genomic structure of the lnc-Rewind locus are shown in the magnified box. **B)** UCSC visualization showing the chromosome position and the genomic coordinates of hs_lnc-Rewind (red box) in the hg19 human genome. The reads coverage from a RNA-Seq experiment performed in proliferating (MB) myoblast (Legnini et al., 2017; GSE70389) are shown below. **C)** Real-time RT-PCR (qRT-PCR) quantification of lnc-Rewind in myoblast (C_2_C_12_) and muscle satellite cells (MuSCs) in growth (GM) and differentiated (DM) conditions. Data represent the mean ± SEM of 3 biological replicates and were normalised on GAPDH mRNA. **D)** Semiquantitative RT-PCR (sqRT-PCR) quantification of lnc-Rewind in cytoplasmic (Cyt) and nuclear (Nuc) fractions from proliferating C2C12 and MuSCs. The quality of fractionation was tested with mature (GAPDH) and precursor (pre-GAPDH) RNAs. **E)** RNA-FISH analysis for lnc-Rewind RNA (red) in proliferating MuCS cell culture. Autofluorescence (grey) is shown with false color to visualize the cell body. DAPI, 4’,6-diamidino-2-phenylindole (blue); Scale bar: 25 µm. **F)** Digital magnification and 3D-visualization of the square insert in panel E. Asterisks indicate lnc-Rewind RNA signals inside the nuclear volume. DAPI, 4’,6-diamidino-2-phenylindole (blue). Data information: *P<0.05, paired Student’s t-test.

To pinpoint possible roles of the murine transcript in myogenic differentiation we first assessed lnc-Rewind expression and subcellular localization both in C_2_C_12_ and FACS-isolated Muscle Stem Cells (MuSCs) (Figure S1C) grown under proliferative and differentiating conditions. Proper myogenic differentiation was confirmed by the expression of late muscle-specific genes such as myogenin (*MyoG*) and muscle creatin kinase (*MCK*) (Figure S1D). Quantitative qRT-PCR analysis revealed that lnc-Rewind expression is high in proliferating (GM) C_2_C_12_ myoblasts and MuSCs and significantly decreased in fully differentiated (DM) cells (Figure 1C). Subcellular fractionation of cytoplasmic (Cyt) and nuclear (Nuc) fractions (Figure 1D) showed that the majority of the noncoding transcript localises in the nuclear compartment of both proliferating myoblasts and MuSCs. Accordingly, RNA fluorescence *in situ* hybridization (FISH) experiments performed both in MuSCs (Figures 1E-1F and S1E) and C_2_C_12_ (Figure S1F) cells confirmed the nuclear localization of lnc-Rewind and further revealed its specific enrichment to discrete chromatin foci. Overall these results point towards a role for lnc-Rewind in chromatin-based processes and suggest its possible involvement in the epigenetic regulation of MuSCs homeostasis.

### Lnc-Rewind regulates muscle system processes and MuSCs expansion

To gain insights into the functional role of lnc-Rewind and to identify the molecular pathways involved in the regulation of MuSC, we performed a global transcriptional profiling on satellite cells treated either with a mix of three different LNA GapmeRs against lnc-Rewind (GAP-REW) or with a scramble control (GAP-SCR) (n=4) (Figure S2A, upper panel). Under these conditions, we obtained ∼70% reduction of lnc-Rewind expression (Figure S2A, lower panel), which led to the identification of a set of 1088 Differentially Expressed Genes (DEGs) (p<0.05, GAP-SCR *vs* GAP-REW). Of these, 332 were upregulated and 756 downregulated in GAP-REW as compared to the scramble condition (Figure 2A and **Table S1**). The PCA clustering analysis showed that the two GAP-SCR and GAP-REW experimental groups displayed a clear different pattern of gene expression since they occupy different regions of the PCA plot (Figure S2B). The prioritized DEGs list was then subjected to Gene ontology (GO) term enrichment analysis (Biological process) to define functional clusters. It emerged that DEGs were mostly associated with muscle cell physiology (skeletal muscle contraction, P-value= 4.23E-6) (Figure 2B and Figure S2C). Of note, the analysis of lnc-Rewind generated miRNAs (let7b and let7c2) revealed that although lnc-Rewind depletion results on their concomitant downregulation (Figure S2D), none of the up-regulated transcripts that are also putative let7b and let7c2 targets (∼1% of the up-regulated genes) (Figure S2E), belong to any of the GO enriched categories. This result emphasizes a specific and miRNA-independent role for lnc-Rewind. Both “Muscle system process” (P-value= 3.45E) and “striated muscle contraction” (P-value= 4.63E-7) were the most significantly enriched GO terms (Figure S2C). Of note, genes encoding for different proteins involved in muscle contraction, such as myosins (i.e. *Myh8*, *Myh3*, *Myll*) and troponins (i.e. *Tnntl*, *Tnnt2*) (Figure 2C) were downregulated. Accordingly, lnc-Rewind depleted MuSCs express lower levels of MyHC proteins (Figure 2D). Morphological evaluation highlighted a decreased number of MuSCs after lnc-Rewind depletion (Figure S2F), suggesting a primary defect in MuSCs proliferation. This led us to hypothesize that the defects in myogenic capacity (Figure 2C-2D and S2F) might result by a decreased cell density that normally leads to a lower rate of myogenic differentiation. In line with this hypothesis, EdU incorporation experiments revealed a striking reduction of proliferating MuSCs upon depletion of Lnc-Rewind (Figure 2E). Accordingly, the *Ccnd3* gene encoding for Cyclin D3, a cyclin specifically involved in promoting transition from G1 to S phase, was significantly downregulated in lnc-Rewind-depleted MuSCs at both transcript (**Table S1**) and protein levels (Figure 2D). In further support of this, MuSCs on single myofibers cultured for 96 hours gave rise to a decreased percentage of proliferating (Pax7^+^/Ki67^+^) progeny upon *lnc-Rewind* downregulation (Figure 2F). Moreover, quantification of the number of Pax7^+^-derived clusters (composed of more than 2 nuclei), pairs (composed of 2 nuclei) or single MuSCs within each myofiber revealed a reduction of activated pairs and clusters upon lnc-Rewind depletion (Figure 2G), suggesting a role for the lncRNA in sustaining MuSCs activation and expansion.

**Figure 2:**
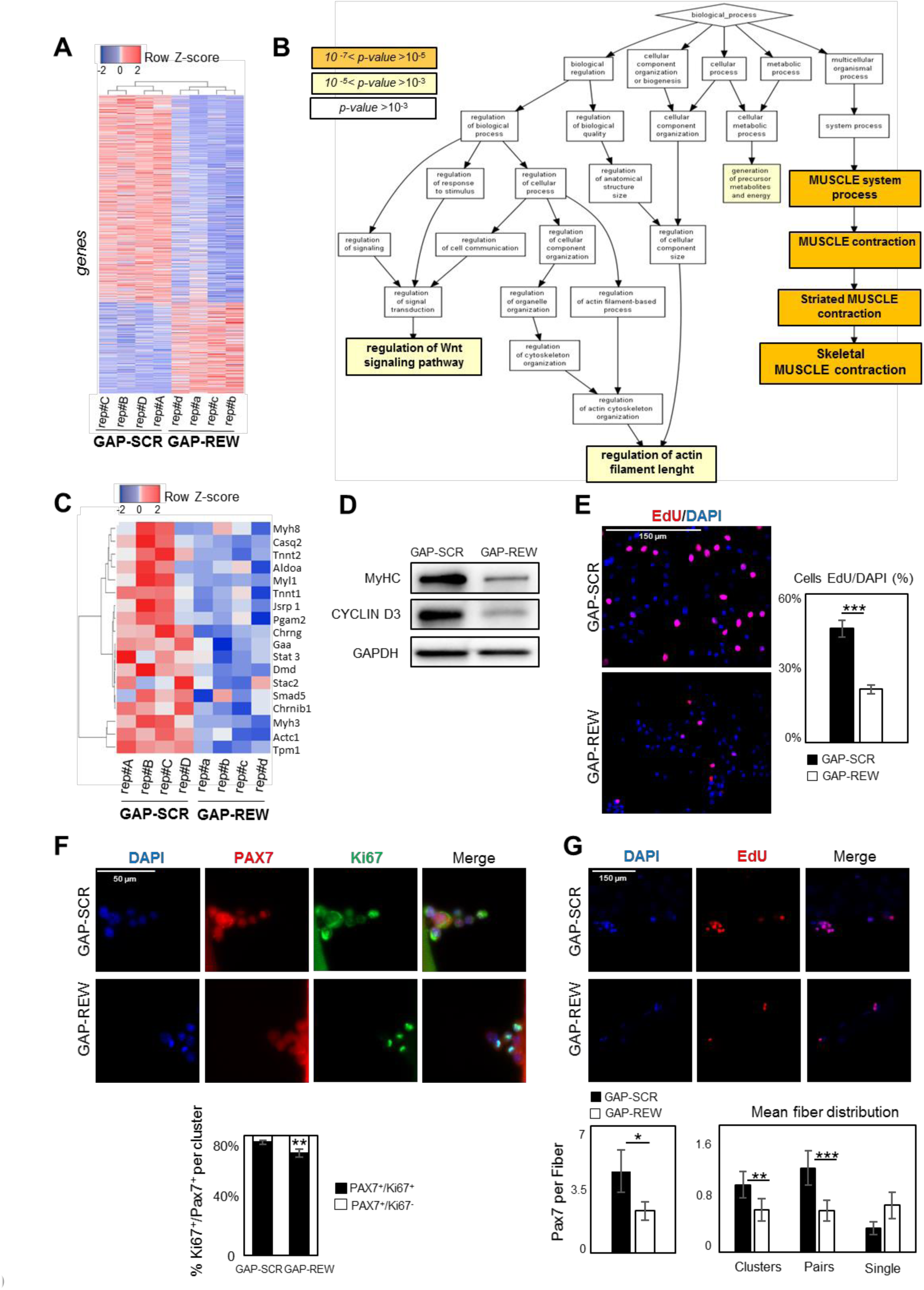
Lnc-Rewind regulates muscle system processes and MuSCs expansion. **A)** Heatmap represents hierarchical clustering performed on the common genes differentially expressed upon lnc-Rewind depletion in MuSC. The analysis was performed using Heatmapper webserver tool (Babicki et al., 2016). The expression levels for each gene correspond to their Z-score values. **B)** Gene Ontology (GO) enrichment analysis performed by GORILLA (Eden et al., 2009) in Biological Process for genes differentially expressed in response to lnc-Rewind depletion in MuSC. **C)** Heatmap represents hierarchical clustering performed on the common genes differentially expressed and belonging to the Skeletal Muscle contraction GO category (GO:0003009). The analysis was performed using the heatmapper webserver tool (Babicki et al., 2016). The expression levels for each gene correspond to their Z-score values. **D)** Western blot analysis performed on protein extracts from MuSCs treated with GAP-SCR or GAP-REW. **E)** Representative images of MuSCs treated with GAP-SCR and GAP-REW, incubated for 6 hours with EdU (red). Nuclei were visualized with DAPI (blu) (left panel) and the histogram shows the percentage of EdU positive cells on the total of DAPI positive cells. Data are graphed as mean ± SEM of 13 images with a total of more then 1000 cells per condition; N=1 mouse (right panel). **F)** Representative images of single muscle fibers treated with GAP-SCR or GAP-REW from Wt mice after 96 hours in culture, stained for Pax7 (red) and Ki67 (green). Nuclei were visualized with DAPI (blue) (upper panel); the histogram shows the percentage of Pax7^+^/Ki67^-^ cells on the total of Pax7^+^/Ki67^+^ cells per cluster (40 clusters per condition; n=5 mice) (lower panel). Data represent the mean ± SEM. **G)** Representative images of single muscle fibers treated with GAP-SCR or GAP-REW from Wt mice after 96 hours in culture, incubated for 24 hours with EdU (red). Nuclei were visualized with DAPI (blu) (upper panel); histograms represent the mean of clusters (n>2), pairs (n=2) and single cells PAX7^+^ (n=1) per fiber. Data represent the mean ± SEM of 80 fibers per condition; n=5 mice (lower panel, right). The histogram (lower panel, left) shows the number of Pax7+ cells per fiber (50 fibers per condition; n=5 mice). Data information: statistical analysis was performed using unpaired Student’s t-test: *P<0.05, **P<0.01, ***P<0.001.

### *Lnc-Rewind* and *Wnt7b* genes display opposite pattern of expression

Together with the muscle-specific genes, the “regulation of Wnt signaling pathway” term caught our attention as it was represented by a significant subset of trancripts (P-value= 5.44E-4, Figure S2C). Among them we found *Wnt7b*, which expression was found upregulated at both transcript (**Table S1**) and protein level (Figure S2G), suggesting a role for *lnc-Rewind* as a repressor of *Wnt7b* expression. Intriguingly, *Wnt7b* transcriptional locus localizes only100 kb upstream *lnc-Rewind* gene (Figure 3A). Their anticorrelated expression together with the lncRNA chromatin enrichment (Figure 1D) and the proximity between the two transcription loci, led us to hypothesize that lnc-Rewind might have a direct, *in cis*-regulatory role on *Wnt7b* transcription. A first clue in favour of such hypothesis came from FANTOM5 Cap Analysis Gene Expression (CAGE) profiles of mouse samples, available on ZENBU genome browser (Noguchi et al., 2017). Indeed, the inspection of the Transcriptional Start Site (TSS) usage among all the available samples (n=1196) revealed a distinctive anti-correlated expression of murine lnc-Rewind and Wnt7b transcripts (Figure 3B, left panels). In contrast, the expression of the other lnc-Rewind-neighbouring genes, such as Cdpf1 and Atxn10, displayed no specific expression correlation (Figure 3B, right panels). Of note, among the different genes analysed in proximity to *Lnc-Rewind* gene, Wnt7b was the only gene significantly upregulated upon the lncRNA depletion (Figure 3C-D) with a concomitant increase in WNTb protein levels (Figure S2G). Although the function of WNT7b has been studied in different cell types and developmental processes (Chen et al., 2014; Wang et al., 2005), the involvement of this ligand in muscle biology is still rather unexplored. The fact that in MuSCs, *Wnt7b* gene is mantained at very low levels and that it is latest Wnt ligand induced after muscle injury (Polesskaya et al., 2003), suggests that the role of WNT7b in muscle might be limited to late stages of myogenic differentiation and implies the necessity to keep it repressed in MuSCs. Our results candidates Lnc-Rewind as a regulator of *Wnt7b* repression in muscle and prompted us towards the study of the underlying mechanism by which this regulation occurs.

**Figure 3:**
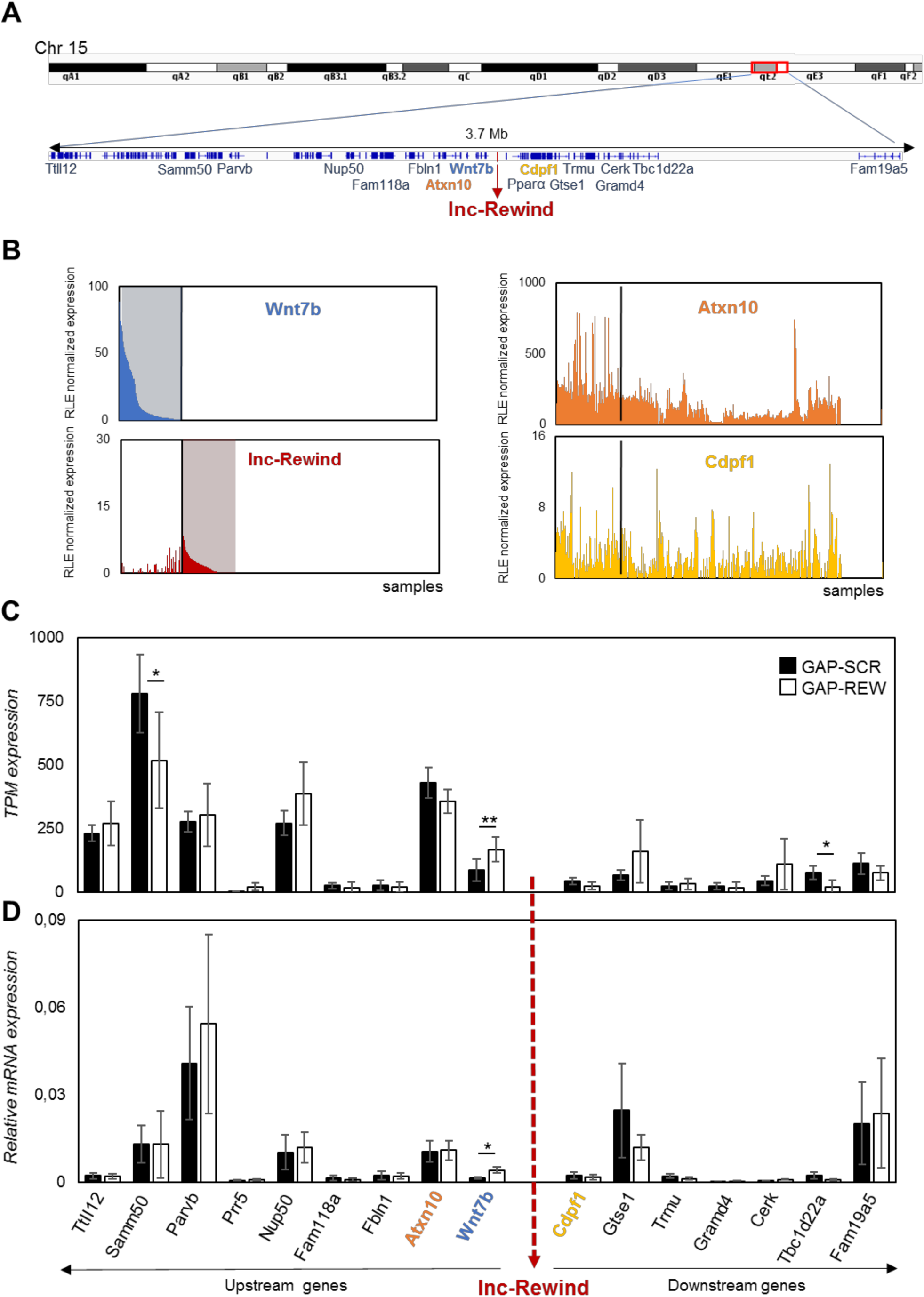
*Lnc-Rewind* and *Wnt7b* genes display opposite pattern of expression. **A)** Schematic UCSC visualization showing the chromosome position and the lnc-Rewind neighbouring genes. **B)** TSS usage analysis of Wnt7b, lnc-Rewind, Atxn10 and Cdpf1 performed by using FANTOM5 (Phase1 and 2) CAGE datasets. Each bar represents the relative logarithmic expression (rle) of the Tag Per Million (TPM) values of the TSS of each gene in one sample (1196 samples). The order of the samples is the same in each of the histograms. **C)** TPM expression (from RNA-seq analysis) of lnc-Rewind neighbouring genes in GAP-SCR *versus* GAP-REW treated MuSCs. TPM expression values are presented as an average±SD of 4 biological replicates. **D)** qRT-PCR quantification of lnc-Rewind neighbouring genes in GAP-SCR *versus* GAP-REW treated MuSCs. Data were normalized to GAPDH mRNA and represent the average ±SEM of 4 biological replicates. Data information: *P<0.05, **P<0.01, paired Student’s t-test.

### Lnc-Rewind directly interacts with the methyltransferase G9a and mediates specific *in-cis* repression of *Wnt7b* in MuSCs

The above results suggest that Lnc-Rewind exerts a repressive role on Wnt7b expression, Several works accumulated so far indicate that most of the *cis*-acting chromatin-associated lncRNAs function occurs by recruiting and guiding chromatin modifiers to target genes (Batista and Chang, 2013; Guttman and Rinn, 2012; Khalil et al., 2009; Rinn and Chang, 2012). In light of our data showing a negative correlation of *Wnt7b* expression by Lnc-Rewind, we focused our attention on the two most known repressive lysine methyltransferases, EZH2 and G9a, which catalyse the deposition of H3K27me3 and H3K9me1/2 on target genes, respectively (Mozzetta et al., 2015). Both EZH2 (Caretti et al., 2004) and G9a (Ling et al., 2012) are mostly expressed in proliferating MuSCs and become downregulated during muscle differentiation (Figure S3A), similarly to Lnc-Rewind. Moreover, both Ezh2 (Rinn et al., 2007; Zhao et al., 2008) and G9a (Nagano et al., 2008; Pandey et al., 2008) have been previously reported to be recruited to specific genomic loci trough the interaction with different lncRNAs. Thus, we hypothesized that in proliferating myoblasts Lnc-Rewind might interact with Ezh2 and/or G9a repressive complexes to tether them on *Wnt7b* genomic locus. Therefore, we performed RNA immunoprecipitation (RIP) analysis in proliferating C_2_C_12_ cells using antibodies against G9a and Ezh2. We observed that, although both G9a and Ezh2 were successfully immunoprecipitated (Figure 4A, upper panel), Lnc-Rewind transcript was efficiently retrieved only in the G9a native RNA immunoprecipitation (IP), while it was almost undetectable in Ezh2 IP (Figure 4A, lower panel). The specificity of the interaction between Lnc-Rewind with G9a, was further confirmed through Crosslinked Immunoprecipitation (CLIP) assay (Figure 4B). Indeed, we found that lnc-Rewind was specifically enriched in the G9a immunoprecipitation if compared to the GAPDH negative control. To note, the levels of RNA obtained in the specific IP fraction were the same of the Kcnq1ot1 lncRNA which was previously demonstrated to be physically associated with G9a (Pandey et al., 2008).

**Figure 4:**
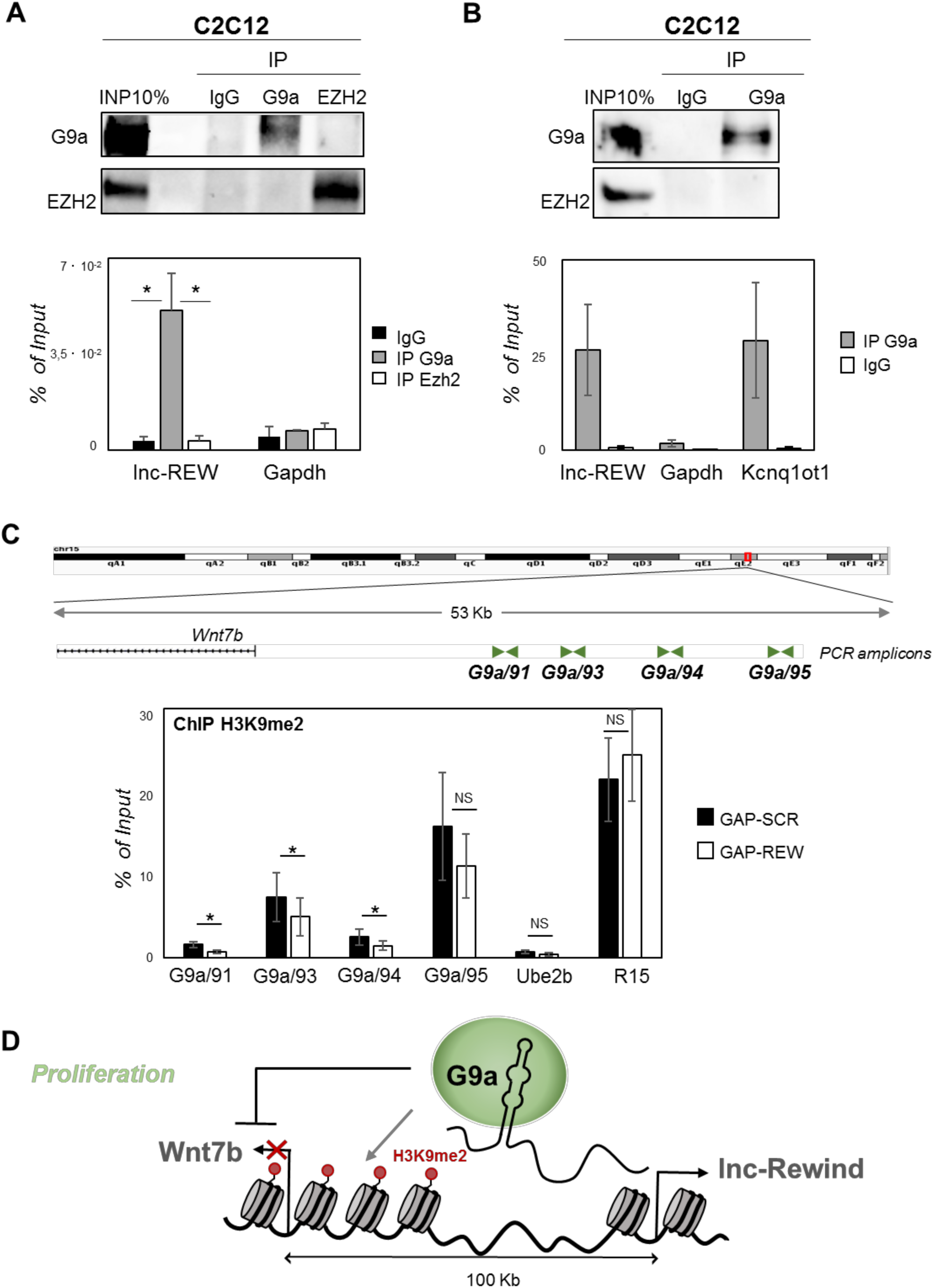
Lnc-Rewind directly interacts with the methyltransferase G9a and mediates specific *in-cis* repression of *Wnt7b*. **A)** G9A and EZH2 native RNA immunoprecipitation (RIP) performed on nuclear extracts of C_2_C_12_ proliferating myoblasts. Data represent mean ±SEM of 4 biological replicates. Western blot analysis of G9A and EZH2 (upper panel) and qRT-PCR quantification of lnc-Rewind recovery (lower panel) are shown. Gapdh RNA serves as negative control. **B)** G9a Cross-linked RNA immunoprecipitation (CLIP) performed on nuclear extracts of C_2_C_12_ proliferating myoblasts. Data represent mean ±SEM of 3 biological replicates. Western blot analysis of G9A (upper panel) and qRT-PCR quantification of lnc-Rewind recovery (lower panel) are shown. Gapdh RNA and EZH2 protein serve as negative controls; Kcnq1ot RNA was used as positive control. **C)** A zoom in into the genomic regions upstream *Wnt7b* gene. Amplicons used to check the H3K9me2 epigenetic mark are shown. Chromatin immunoprecipitation (ChIP) of H3K9me2, performed in Wt MuSCs upon lnc-Rewind downregulation; MuSCs transfected with scrambled (GAP-SCR) gapmers have been used as control. Enrichment is represented as percentage of input DNA (% input). Ube2b and R15 genomic regions were used as negative and positive control, respectively. The graph shows the averaged value derived from n=6 independent experiments. **D)** Proposed model for the murine lnc-Rewind mode of action in muscle cells. In proliferating myoblasts, lnc-Rewind is expressed and guide the silencing methyltransferase G9a on *Wnt7b* promoter to repress its transcription. Data information: *P<0.05, paired Student’s t-test.

In order to test whether Lnc-Rewind/G9a interaction has a role for the deposition of H3K9me2 mark on *Wnt7b* containing regions, we performed a chromatin immunoprecipitation (ChIP) assay for H3K9me2 in scramble and Lnc-Rewind gapmer transfected MuSCs. We analysed different regions upstream *Wnt7b* TSS (Figure 4C, upper panel) and quantification of H3K9me2 enrichment led to the identification of three regions (named G9a/91, G9a/93 and G9a/94) in which H3K9me2 levels decreased upon Lnc-Rewind knockdown, compared to the SCR control (Figure 4C, lower panel). Taken together, these results support a mechanism of action through which in MuSCs Lnc-Rewind represses *Wnt7b* gene transcription by mediating the specific G9a-dependent H3K9me2 deposition on its locus (Figure 4D).

## DISCUSSION

Wnt signalling represents one of the pathways that has a major role in myogenesis as it is essential for proper MuSCs self-renewal and differentiation (von Maltzahn et al., 2012). A correct timing of Wnt signalling activation is crucial to obtain proper tissue repair and its aberrant activity causes a wide range of pathologies (Nusse and Clevers, 2017) Thus, it is not surprising that Wnt pathway is under a strict positive and negative multi-layered regulation. Recent studies show that long noncoding RNAs can modulate Wnt pathway by affecting gene expression through different mechanisms, from transcriptional to post-transcriptional level (Ong et al., 2017; Shen et al., 2017; Zarkou et al., 2018) For example, lncRNAs were found interacting with transcription factors (Di Cecilia et al. 2016) and chromatin modifiers (Hu et al., 2015) altering Wnt signaling pathway in different tissues and in cancer. In this study we discovered that lncRNAs-mediated regulation plays a role in modulating Wnt pathway in muscle stem cells. Taking advantage of a newly discovered atlas of not annotated muscle-specific lncRNAs (Ballarino et al., 2015), we decided to focus our attention on Lnc-Rewind for the following reasons: i) an orthologue transcript, hs_lnc-Rewind, with high level of sequence identity is expressed by human myoblasts (Figure 1B **and** S1A-B); ii) lnc-Rewind is associated and retained to chromatin (Figue 1D-F); iii) it is in genomic proximity to *Wnt7b* locus (Figure 3A) and iv) its knock-down induces upregulation of Wnt7b in MuSCs (Figure 3C and S2G). These observations led us hypothesize a Lnc-Rewind-mediated *in-cis* repression for *Wnt7b* gene. Of note, by analyzing the expression of surrounding genes within Lnc-Rewind genomic region, *Wnt7b* is the only gene to be significantly up-regulated by Lnc-Rewind depletion (Figure 3C-D). Moreover, we show that Lnc-Rewind and *Wnt7b* expression are anti-correlated in other cell types (Figure 3B), suggesting that the repressive action of Lnc-Rewind on *Wnt7b* locus could have a more general role, not only in satellite cells.

It is well-accepted that the majority of the *cis*-acting chromatin-associated lncRNAs function by recruiting and targeting chromatin modifiers to specific genes (Batista and Chang, 2013; Guttman and Rinn, 2012; Khalil et al., 2009; Rinn and Chang, 2012). In light of the evidence that Lnc-Rewind promotes *Wnt7b* repression, we decided to investigate the involvement of the two major repressive histone modifiers, Ezh2 and G9a, in mediating such Lnc-Rewind-dependent silencing. Notably, by performing two different biochemical approaches (RIP and CLIP), we found that only G9a, specifically interacts with Lnc-Rewind (Figure 4A-B). This strongly suggests that this histone H3K9 lysine methyltransferase (KMT) might be specifically tethered on *Wnt7b* genomic locus by Lnc-Rewind to mediate its transcriptional repression. In support to this idea, through H3K9me2 ChIP experiments here we provide evidence that different regions upstream *Wnt7b* locus are enriched in the G9a-deposited histone mark in proliferating MuSCs and that H3K9me2 levels on these regions significantly decrease upon lnc-Rewind knockdown (Figure 4C). Finally, expression profiling of Lnc-Rewind-depleted MuSCs strongly confirmed a functional role for this novel lncRNA as a modulator of Wnt signalling (Figure 2B), myogenic capacity (Figure 2B-C-D and S2F) and MuSCs expansion (Figure 2E-F-G).

A correct timing and magnitude of Wnt signalling activation is essential to maintain functionl MuSCs. For instance, it has been shown that low β-catenin activity is fundamental during early phases of muscle regeneration to allow MuSCs activation and subsequent differentiation (Figeac and Zammit, 2015; Parisi et al., 2015). Accordingly, satellite cells isolated from mice with constitutive active Wnt/β-catenin signalling display an early growth arrest and premature differentiation (Rudolf et al., 2016). All these data point out that the cell environment, together with the timing and the proper activation of Wnt signaling pathway is foundamental for muscle homeostasis. Here we show that aberrant, cell-autonomous activation of Wnt7b expression upon Lnc-Rewind depletion causes defects in MuSCs expansion and activation (Figure 2E-F-G), emphasizing that Lnc-Rewind-mediated repression of *Wnt7b* is key in ensuring MuSCs activity. This is nicely supported by a previous study demonstrating that, in the cytoplasm, a Wnt7b/lncRNA circuitry controls the proliferation of C_2_C_12_ myoblasts. Specifically, Lu and colleagues demonstrated that the YY1-associated muscle lincRNA (Yam-1) inhibits skeletal myogenesis through modulation of miR-715 expression, which in turn targets Wnt7b mRNA (Lu et al., 2013). In sum, our results support a mechanism of action through which Lnc-Rewind mediates G9a recruitment on *Wnt7b* locus to silence it through the deposition of the repressive mark H3K9me2 during the proliferation phase of satellite cells (Figure 4D), thus providing unique insights on the contribution of nuclear-enriched lncRNAs in myogenesis.

## DATA AND SOFTWARE AVAILABILITY

Data that support the findings of this study have been deposited in GEO with the primary accession code GSE141396

https://www.ncbi.nlm.nih.gov/geo/query/acc.cgi?acc=GSE141396; <u>GEO reviewer token: mlqfamksbrirvip</u>

## SUPPLEMENTAL INFORMATION

Table S1.

## ACKNOWLEDGMENTS

This work was partially supported by grants from Sapienza University (prot. RM11715C7C8176C1 and RM11916B7A39DCE5) and FFABR Anvur (2017) to M.B.. C.M. laboratory is funded by Italian Ministry of University and Research (SIR, Scientific Independence of Young Researcher n. RBSI14QMG0), Italian Association for Cancer Research (AIRC; MyFIRST grant n.18993), AFM-Telethon (#22489) and the CNCCS (Collection of National Chemical Compounds and Screening Center), LIFE2020-Regione Lazio.

## AUTHOR CONTRIBUTIONS

C.M. and M.B. conceived, supervised the project and wrote the manuscript. A.C. performed the cellular and molecular analyses in C_2_C_12_ cells, RNA-seq analysis, TSS usage, Gene Onthologies studies, ChIP experiments and the statistical analyses; M.M. performed the cellular and molecular experiments carried out on MuSCs and myofibers, RIP and CLIP on C_2_C_12_; B.B. contributed to the cellular experiments on MuSCs and myofibers. G.P. perfomed FACS isolation of MuSCs. T.S. performed RNA-FISH; G.B. contributed to the molecular analyses performed in C_2_C_12_ and MuSCs cells; Al. Col. performed the bioinformatic analysis of the human myoblast RNA-seq. All authors discussed results, reviewed and edited the manuscript.

## METHODS

### Ethics statement and animal procedures

For the experiments described in this study, C57/BL10 wild-type mice were used and differences which were observed in both male and female mice were included in experiments. Animals were treated in respect to housing, nutrition and care according to the guidelines of Good laboratory Practice (GLP). All experimental protocols were approved and conformed to the regulatory standards. All animals were kept in a temperature of 22°C ± 3°C with a humidity between 50% and 60%, in animal cages with at least 5 animals.

### Cell preparation and FACS sorting

Cell isolation and labeling was essentially performed as described in (Mozzetta, 2016). Isolation of cells from two months old C57/Bl10 WT mice was performed as follows: briefly, whole lower hindlimb muscles were carefully isolated, minced and digested in PBS (Sigma) supplemented with 2.4 U/ml Dispase II (Roche), 100 μg/ml Collagenase A (Roche), 50 mM CaCl2, 1 M MgCl2, 10 mg/ml DNase I (Roche) for 1 hour at 37 °C under agitation. Muscle slurries were passed 10 times through a 20G syringe (BD Bioscience). Cell suspension was obtained after three successive cycles of straining and washing in Washing Buffer consisting of HBSS containing 0.2% BSA (Sigma Aldrich), 1% penicillin-streptomycin. Cells were incubated with primary antibodies CD31-PE (MiltenyBiotec, 130111540), CD45-PE (MiltenyBiotec, 139110797), Ter119-PE (MiltenyBiotec, 130112909) 1:25; Sca1-FITC (MiltenyBiotec, 130116490) 1:25 e α7Integrin-APCVio770 (MiltenyBiotec, 130102719) 1:20 for 45 min on ice diluted in HBSS containing 0.2% BSA, 1% penicillin-streptomycin and 1% DNAse I. The suspension was finally washed and resuspended in PBS containing 2% Fetal Bovine Serum (FBS) and EDTA 0.5 μM. Cells were sorted using a FACSAriaIII (Becton Dickinson, BD Biosciences) equipped with 488nm, 561nm and 633nm laser and FACSDiva software (BD Biosciences version 6.1.3). Data were analyzed using a FlowJo software (Tree Star, version 9.3.2). Briefly, cells were first gated based on morphology using forward versus side scatter area parameter (FSC-A versus SSC-A) strategy followed by doublets exclusion with morphology parameter area versus width (A versus W). Muscle satellite cells (MuSCs) were isolated as Ter119_neg_/CD45_neg_/CD31_neg_/a7-integrin_pos_/Sca1_neg_ cells (see Figure S1C, left and right panels). To reduce stress, cells were isolated in gentle conditions using a ceramic nozzle of size 100µm, a low sheath pressure of 19.84 pound-force per square inch (psi) that maintain the sample pressure at 18.96 psi and a maximum acquisition rate of 3000 events/s. Cells were collected in 5 polypropylene tubes. Following isolation, an aliquot of the sorted cells was evaluated for purity at the same instrument resulting in an enrichment >98-99% (see Figure S1C right panel).

### Cell culture conditions and transfection

Freshly sorted cells were plated on ECM Gel (Sigma)-coated dishes in Cyto-grow (Resnova) complete medium as a growth medium (GM) and cultured at 37°C and 5% CO2. After 5 days in GM, SCs were exposed, generally for 2 days, to differentiation medium (DM) consisting of DMEM with 5% Horse Serum (HS). Cells were counted on DAPI-stained images using the ImageJ tool “Multi point”. Downregulation of Lnc-Rewind expression was performed at ∼50% confluence by transfection in Lipofectamine 2000 (Invitrogen), with 75nM of LNA GapmeRs (Euroclone), according to manufacturer’s instructions. MuSCs were subjected to two consecutive (overnight) rounds of transfection and harvested 24 hours after second transfection.

GapmeRs were designed against Lnc-Rewind sequence using the Exiqon web tool (http://www.exiqon.com/ls/Pages). Negative Control A (Euroclone) was used as negative (Scramble) control. Sequences are reported in Table S2

For EdU (Invitrogen) detection cells were incubated for 6h and stained using Click-iT^TM^ EdU Alexa Flour^TM^ 594 HCS Assay (Invitrogen) according to the manufacturer’s instructions.

### Single myofibers isolation and immunofluorescence

Single myofibers were isolated from Extensor digitorum longus (EDL) muscles of C57/Bl10 mice and digested in 2mg/ml Collagenase I (Sigma) for 1 h at 37 °C, gently shacked every 10 minutes. Single fibers were obtained gently triturating the digested EDL muscles using a glass pipette in DMEM supplemented with 10% HS (horse serum). The myofibers were manually collected under a dissecting microscope and cultured in DMEM + Pyr with 20% FBS, 2,5 ng/ml FGF (Gibco) and 1% Chick Embryo Extract (CEE) (Life Science Production). Downregulation of Lnc-Rewind expression was performed 4 h after plating with one round of transfection, as described above. The fibers were incubated for 24h with EdU (Invitrogen) and were analyzed 96h after plating.

For EdU detection cells were stained using Click-iT^TM^ EdU Alexa Flour^TM^ 594 HCS Assay (Invitrogen) according to the manufacturer’s instructions.

For the immunofluorescence, the single myofibers were fixed with 4% paraformaldehyde in PBS for 20 min at RT, permeabilized with 0.5% Triton X-100 in PBS, and blocked with 10% FBS in PBS for 1 h at RT. Primary antibodies (Pax7 (DSHB, Pax7-s) 1:10 and Ki67 (Abcam, ab15580) 1:100) were diluted in 10%FBS-PBS and incubated overnight at 4°C. After incubation for 1h with the appropriate secondary antibodies (Alexa Fluor 488 or 594 (Thermo Fisher)), nuclei were counterstained with DAPI (Sigma) and fibers were mounted on cover-glasses. Images were taken with Axio Observer microscope (ZEISS) and processed with ZEN 3.0 (Blue edition) software.

### RNA Imaging

Lnc-Rewind *in situ* hybridization analyses were performed as previously described (Rossi et al., 2019). Briefly, proliferating MuSC and C_2_C_12_ cells were cultured on pre-coated glass coverslips and then fixed in 4% paraformaldehyde/PBS (Electron Microscopy Sciences, Hatfield, PA). RNA hybridization and signal development were carried out using Basescope assay (Advanced Cell Diagnostics, Bio-Techne) and BA-Mm-lnc-Rewind probe-set (ref. 722581) designed to detect three junction regions of Lnc-Rewind (exon/intron1, intron1/exon2, exon/intron2). A probe specific for exon junction exon1/exon2 of human Dlc1 mRNA (BA-HS-DLC1, ref.716041) was used as negative control. The images were collected as Z-stacks (200 nm Z-spacing) with MetaMorph software and post-acquisition processing were performed by FIJI software to the entire image. 3D viewer plugin was used to perform 3D-rendering.

### RNA analyses

Total RNA from myoblasts/myotubes and MuSCs was extracted with TriReagent (Sigma). 0.5-1.0ug of RNA were treated with RNase-free DNase I enzyme (Thermo scientific). Nucleus and Cytoplasmic fractionations was performed using the Paris kit (Ambion, AM1921) according the protocol specifications. RNA was reverse transcribed using the SuperScript VILO Master Mix (Thermo Scientific). Real time quantitative PCRs were performed by using SYBR Green Master mix (Applied Biosystems), according to the manufacturer’s instructions. Relative expression values were normalized to the housekeeping GAPDH transcript.

### Cell lysis and Immunoblot

Total proteins were prepared by resuspending cells in RIPA buffer (50mM Tris-HCl pH 7.4, 150mM NaCl, 0.1% SDS, 0.5% sodium deoxycholate, 1% NP-40, 1mM EDTA, protease and phosphatase inhibitors (Roche)). Protein concentration was determined using a BCA assay (Thermo Fisher Scientific). The cell lysate was denatured at 95 °C for 5 min. The cell lysates were resolved on 4%– 15% TGX gradient gels (Bio-Rad Laboratories) and transferred to nitrocellulose membrane (Amersham). Membranes were blocked with 5% non-fat dried milk in TBS with 0.2% Tween for 1 h at RT, and then incubated with primary antibody overnight at 4°C Primary antibodies used were against: MF20 (MyHC) (DSHB, Mf20-s), G9a (Abcam, ab185050), Ezh2 (Cell Signaling, #3147), Wnt7b (Abcam, ab94915), β-catenin (Santa Cruz biotechnology, sc-7963), Cyclin D3 (Santa Cruz biotechnology, sc-182) and GAPDH (Sigma, G9545). After washing in TBS with 0.2% Tween, membranes were incubated in HRP-conjugated specific secondary antibody (goat anti-rabbit-anti-mouse (IgG-HRP Santa Cruz Biotechnologies)) for 1h at RT. After washing in TBS with 0.2% Tween, blots were developed with Western lightning enhanced chemiluminescence (Thermo Fisher Scientific), the signal detection was performed with the use of ChemiDoc (BioRad).

### RNA Immunoprecipitation (RIP)

C_2_C_12_ cells were washed with PBS and centrifuge at 2000 rpm for 5 minutes and resuspended in Buffer A (20mM Tris HCl pH8.0, 10mM NaCl, 3mM MgCl2, 0.1% NP40, 10% Glicerol, 0.2mM EDTA, 0.4mM PMSF, 1XPIC). In ice for 15 minutes and centrifuge at 2000 rpm for 5 minutes at 4°C to pellet the nuclei. The supernatant was collected as cytoplasmic extract. The pellet was re-washed with buffer A for 3 times. The pellet was resuspended in NT2/Wash buffer (50mM Tris pH7.4, 150mM NaCl, 1mM MgCl2, 0,5% NP40, 20mM EDTA, 1X PIC, 1X PMSF, 1 mM DTT), break with 7mL dounce (thight pestel/B pestel) and centrifuged at 14000 rpm for 30 minutes at 4°C. Protein A/G Magnetic beads (Thermos Scientific) were incubate with IgG, G9a (Abcam, ab185050) and Ezh2 (Cell Signaling, #3147) antibody on rotating wheel ON at 4°C while the nuclear extract was precleared with the beads. 10% of the nuclear extract was collected for INP. The NE was divided in each sample with the coated beads (beads+ Ab) and incubate on rotating wheel ON at 4°C. The beads were washed with NTS buffer and ¼ was collected for protein and ¾ for RNA analyses. RIP qPCR results were represented as percentage of IP/input signal (% input).

### Cross-linking immunoprecipitation (CLIP) assay

CLIP experiments were performed on nuclear extract obtained with some modification of the Rinn et al. protocol (Rinn et al., 2007). Briefly, C_2_C_12_ cells were washed with PBS and crosslinked with UV rays and collected in Buffer A (20mM Tris HCl pH8.0, 10mM NaCl, 3mM MgCl2, 0.1% NP40, 10% Glicerol, 0.2mM EDTA, 0.4mM PMSF, 1XPIC). Cells were centrifuged at 500 xg for 10 minutes and resuspended in Buffer A, the nuclei pellet was resuspended in NP40 lysis buffer (50mM HEPES-KOH, 150Mm KCL, 2Mm EDTA, 1Mm NaF, 0,5% NP40 PH 7.4, 0,5Mm DTT, 100X PIC). Break with 7mL dounce (thight pestel/B pestel) and centrifuged at max speed for 20 minutes at 4°C. The supernatant was quantified with bredford assay. 10% of the nuclear extract was collected for INP. Protein A/G Magnetic beads (Thermos Scientific) were washed with PBS tween 0,02% (1X PBS, 0,02% Tween-20) were incubate in PBS Tween 0,02% with IgG and G9a (Abcam, ab185050) antibody on rotating wheel 1 h at RT. The NE was divided in each sample with the coated beads (beads+ Ab) and incubate on rotating wheel ON at 4°C. The beads where washed 3 times with the NP40 High Salt Buffer 0.5X (25 mM HEPES-KOH pH7.5, 250 mM KCL, 0.025% NP40), 2 times with PNK Buffer (50mM Tris-HCL, 50Mm NaCl, 10mM MgCl2 pH7.5) and resuspended with NP40 lysis buffer. ¼ was taken for protein analysis where was add LSD 5X and DTT 100X, for 5 minutes at 95°C. The rest was used for RNA analysis: the beads were resuspended in proteinase K buffer (200MM Tris-HCL, 300 mM NaCl, 25 mM EDTA, 2%SDS pH 7.5).

### Chromatin Immunoprecipitation (ChIP)

ChIP experiments were performed on chromatin extracts according to the manufacturer’s protocol (MAGnify ChIP; Life Technologies) by O.N. incubation with 3 μg of immobilized anti-H3K9me2 (Abcam ab1220) or rabbit IgG antibodies. A standard curve was generated for each primer pair testing 5-point dilutions of input sample. Fold enrichment was quantified using qRT-PCR (SYBR Green; Qiagen) and calculated as a percentage of input chromatin (% Inp). Data from GAP-SCR *vs* GAP-REW conditions represent the mean of six independent experiments ± SEM. Sequences of the oligonucleotidies used for ChIP analyses are reported in Table S2.

### 3’-end mRNA Sequencing and Bioinformatic analysis

Total RNA was quantified using the Qubit 2.0 fluorimetric Assay (Thermo Fisher Scientific). Libraries were prepared from 100 ng of total RNA using the QuantSeq 3’ mRNA-Seq Library Prep Kit FWD for Illumina (Lexogen GmbH); the quality was assessed by using screen tape High sensitivity DNA D1000 (Agilent Technologies). Sequencing was performed on a NextSeq 500 using a high-output single-end, 75 cycles, v2 Kit (Illumina Inc.). Illumina novaSeq base call (BCL) files were converted in fastq file through bcl2fastq (version v2.20.0.422). Sequence reads were trimmed using trimgalore (v0.4.1) to remove adapter sequences and low-quality end bases (regions with average quality below Phred score 20). Alignment was performed with STAR 2.5.3a (Dobin et al., 2013) on mm10 reference. The expression levels of genes were determined with htseq-count 0.9.1(Anders et al., 2015) by using mm10 Ensembl assembly (release 90). Genes having < 1 Count Per Million (CPM) in at least 4 samples and those with a percentage of multi-mapping reads > 20% were filtered out. Paired t-test was performed to select differentially expressed genes in GAP-REW vs GAP-SCR conditions, setting 0.05 as p-value threshold. Gene ontology term enrichment analyses were performed using GORILLA (Eden et al., 2009) providing the list of genes expressed in at least one of the two conditions as background.

RNA-Seq reads from WT myoblasts cultured in growth medium (Legnini et al., 2017) were downloaded from GEO (GSE70389), preprocessed using Trimmomatic 0.32 software (Bolger et al., 2014) and aligned to human GRCh38 assembly using STAR 2.5.3a. The normalized read coverage tracks (.tdf files) were created and loaded on IGV genome browser (Robinson et al., 2011).

**Table S2:**
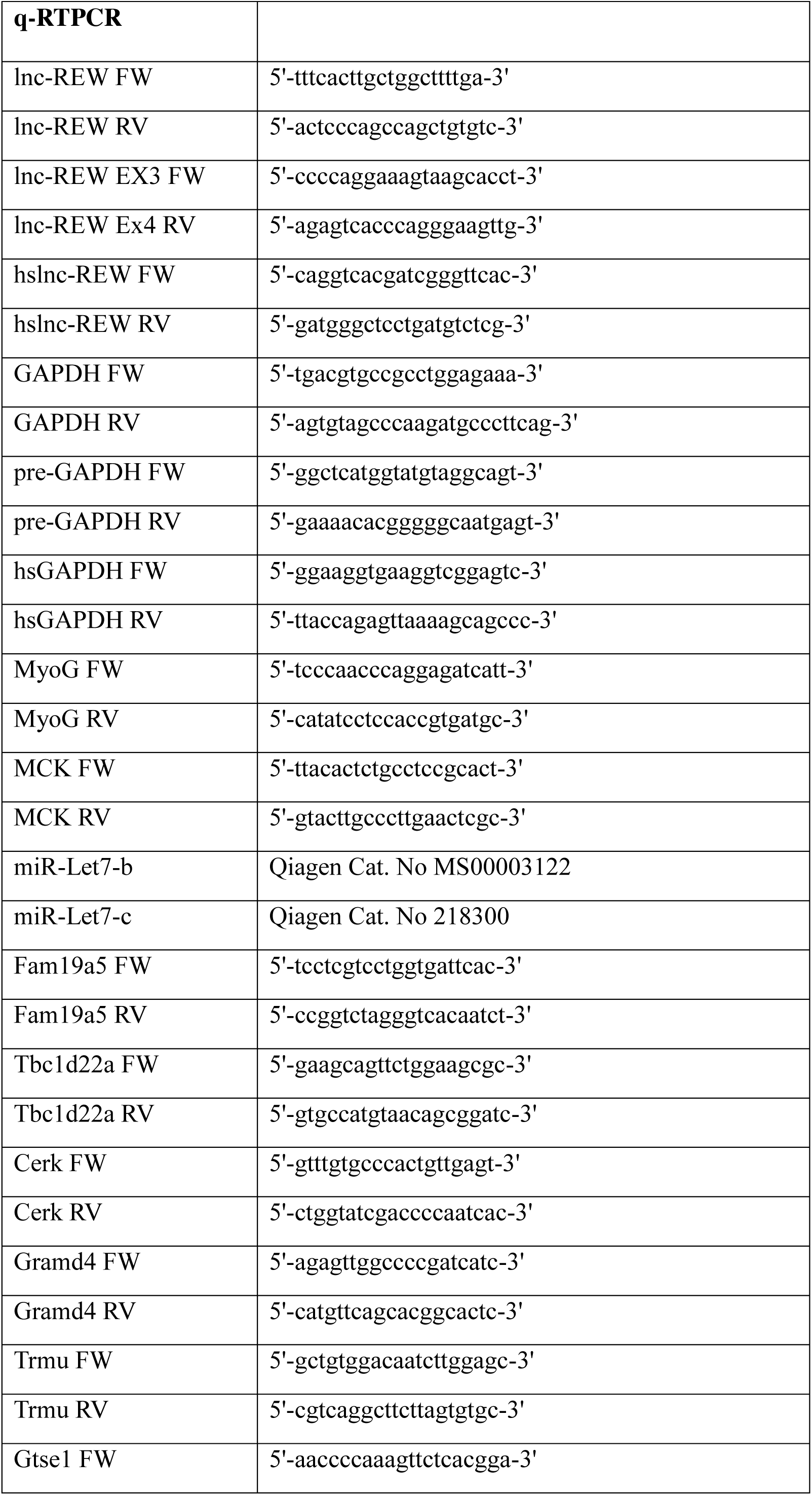
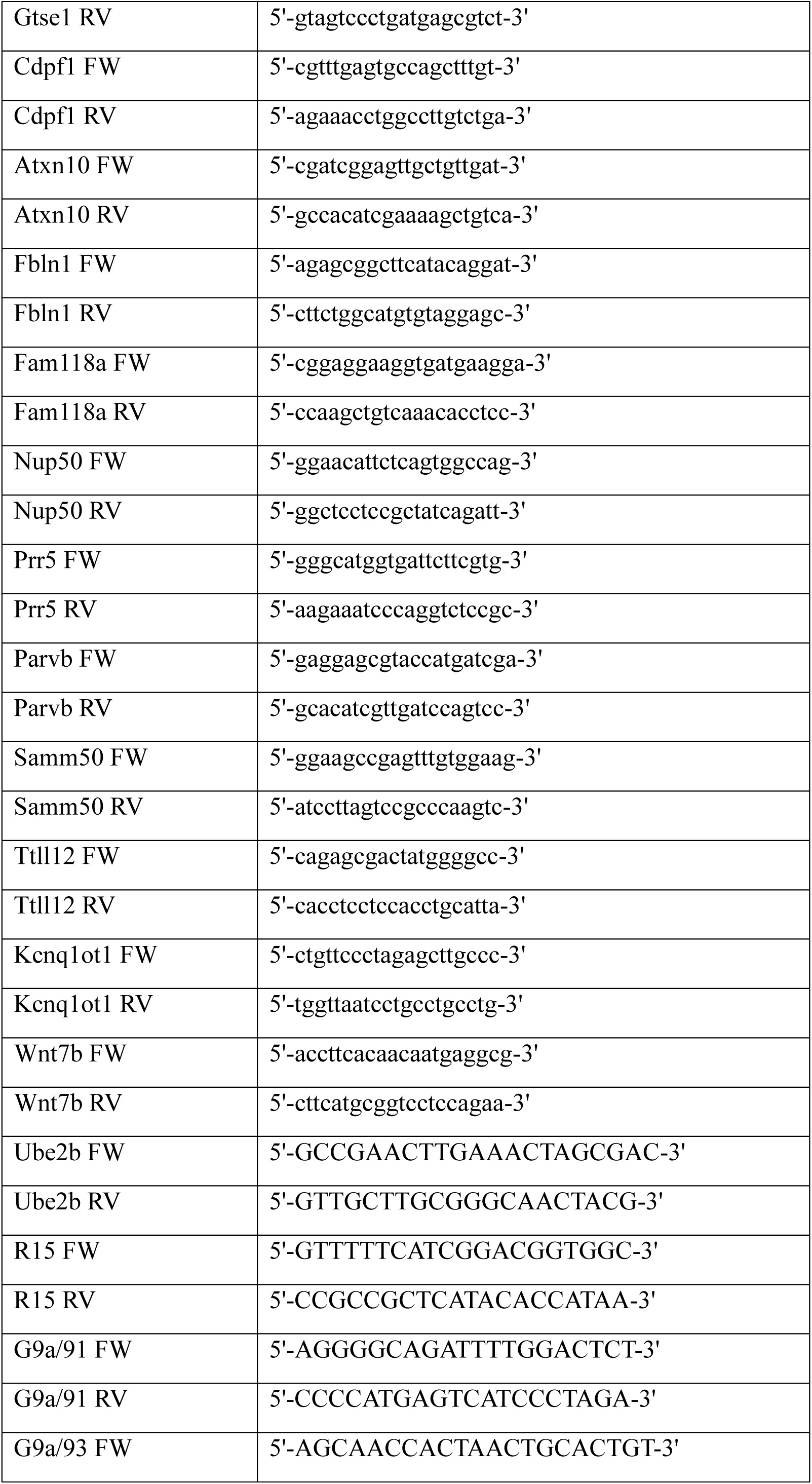
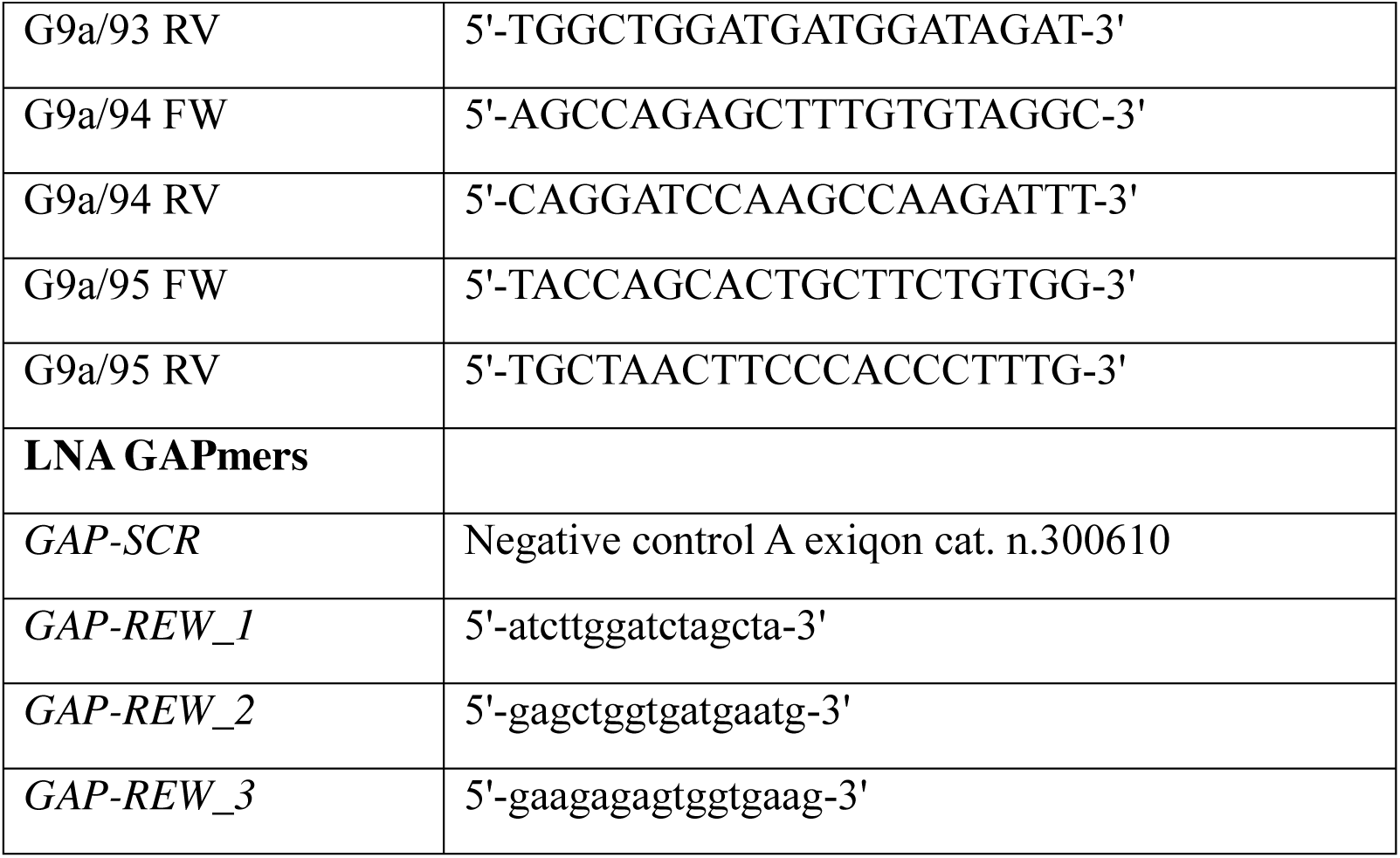
List and sequences of the oligonucleotides and LNA gapmers used.

## SUPPLEMENTAL EXPERIMENTAL PROCEDURES

**Figure S1.**
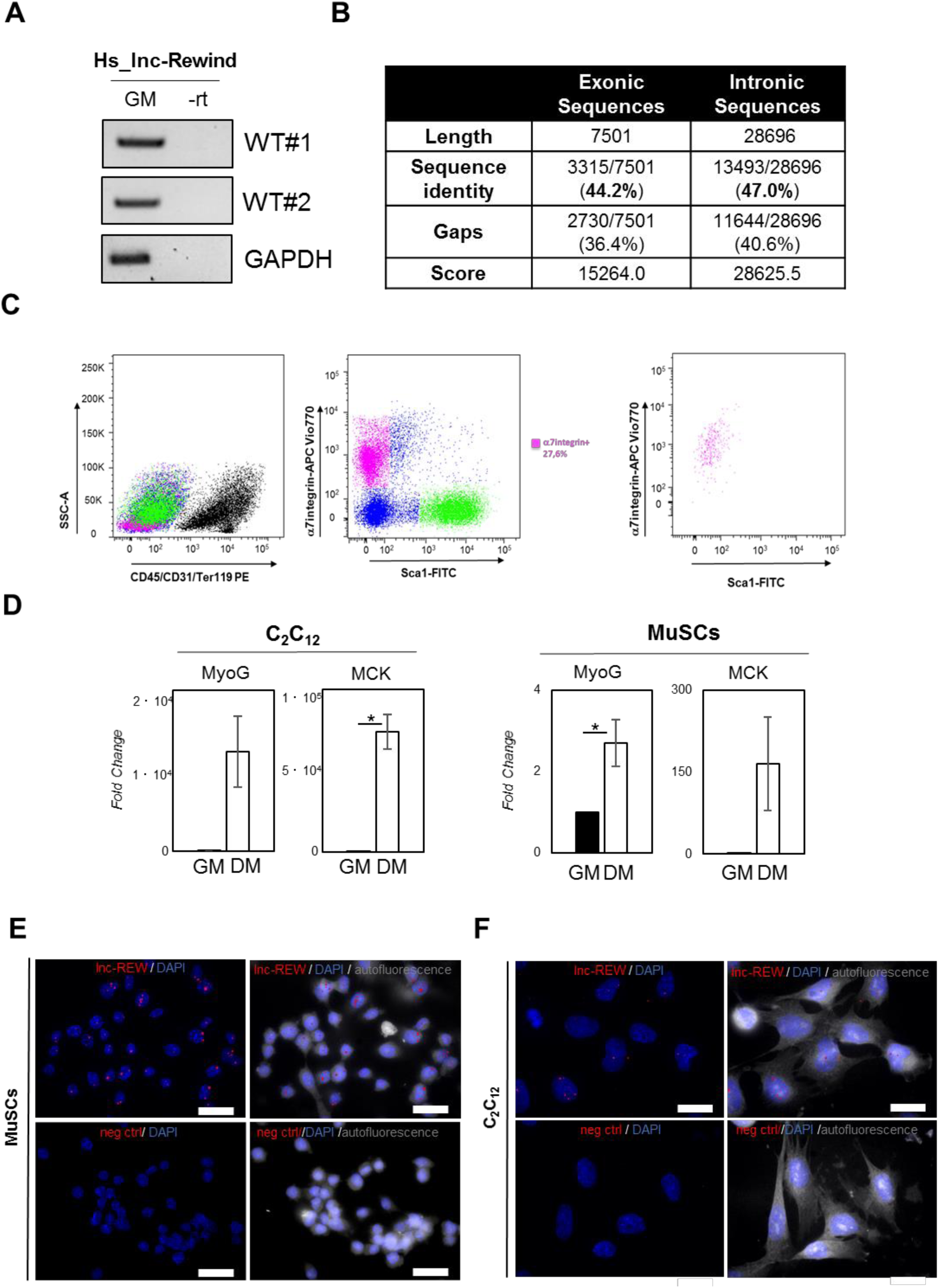
**A)** sqRT-PCR quantification of hs_lnc-Rewind in two different proliferating human myoblasts from healthy donors in proliferating condition. Mature GAPDH was used as an amplification control. **B)** Table represents the values obtained by analysing the local sequence alignment between the human and murine lnc-Rewind transcripts. Data were produced by using the implementation of the Smith-Waterman algorithm available at http://www.ebi.ac.uk/Tools/psa/emboss_water/. **C)** In WT mice, lineage negative CD45/CD31/Ter119 PE cells were sorted for muscle stem cells based on α7integrin expression (APC Vio770 positive cells, magenta) (left and middle panels). The check purity plot shows the pureness of the sample (right panel). **D)** qRT-PCR quantification of MyoG and MCK in C_2_C_12_ and MuSCs in growth and differentiated conditions. Data represent the mean ± SEM of 3 biological replicates and were normalised on Gapdh mRNA. **E)** Representative 60X confocal images of MuSC cell cultures hybridized with probes set specific for lnc-Rewind and for a human mRNA (dlc1). Autofluorescence (grey) is shown with false colour to visualize the cell body. DAPI, 4’,6-diamidino-2-phenylindole (blue); Scale bar: 25 um. **F)** Representative 60X confocal images of C_2_C_12_ cell cultures hybridized with probes set specific for lnc-Rewind and for a human mRNA (dlc1). Autofluorescence (grey) is shown with false colour to visualize the cell body. DAPI, 4’,6-diamidino-2-phenylindole (blue); Scale bar: 25 um. Data information: *P<0.05, paired Student’s t-test

**Figure S2.**
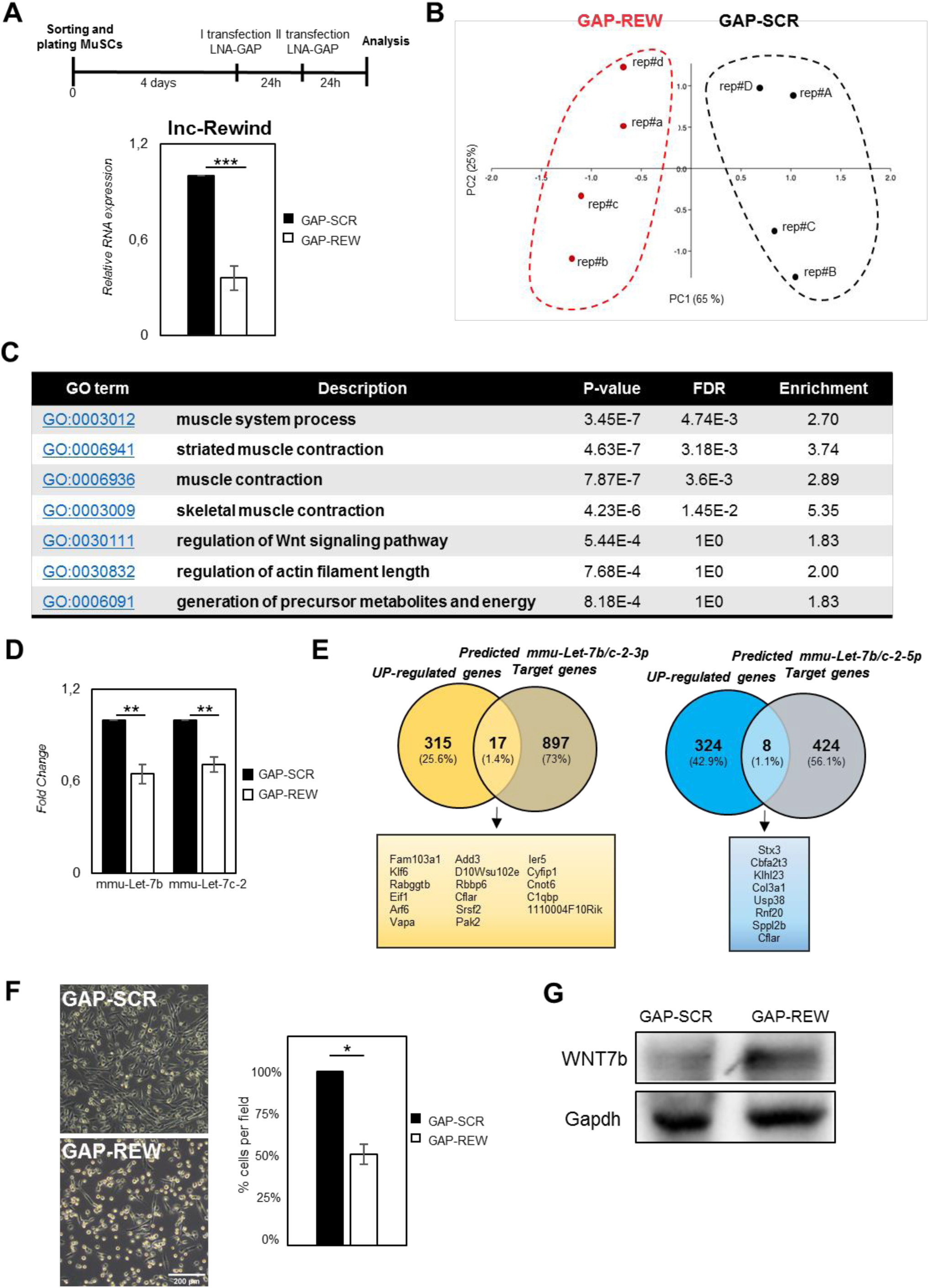
**A)** Schematic representation of the experimental condition of MuSCs treated with GAP-SCR or GAP-REW (upper part). Real-time qRT-PCR quantification of lnc-Rewind in MuSCs treated with GAP-SCR or GAP-REW. Data were normalized to Gapdh mRNA and represent the mean ± SEM of n=4 biological replicates (lower part). **B)** PCA analysis of DEGs upon lnc-Rewind depletion. GAP-SCR and GAP-REW treated samples replicates are shown respectively as red and black dots (left panel). The plot represents each Eingvalue % for the different Principal components (right panel). As shown in both panels, PCA1 and PCA2 represent the top two dimensions of the DEGs which account respectively for 65%, 25% of the total variability. **C)** Table shows the GO terms, their description, the p-value, the FDR (represented in decreasing order of statistical significance) and the enrichment of the different categories showed in Figure 2C. **D)** Real-time qRT-PCR quantification of mmu-Let7-b and mmu-Let7-c-2 in MuSCs treated with GAP-SCR or GAP-REW. Data were normalized to Gapdh mRNA and represent the mean ± SEM of n=3 biological replicates. **E)** Target prediction analysis performed by crossing the lnc-REW up-regulated genes with the mmu-Let7-3p (yellow) and mmu-Let7-5p (Blue) predicted target genes. The predictions were obtained from Targetscan database (Agarwal et al., 2015). **F)** Representative images (left) and cells number quantification (right) of MuSCs treated with GAP-SCR or GAP-REW. Data represent the mean ± SEM from 3 biological replicates. **G)** Western blot analysis performed on protein extracts from MuSCs treated with GAP-SCR or GAP-RE. Data information: *P<0.05, **P<0.01, ***P<0.001, paired Student’s t-test

**Figure S3.**
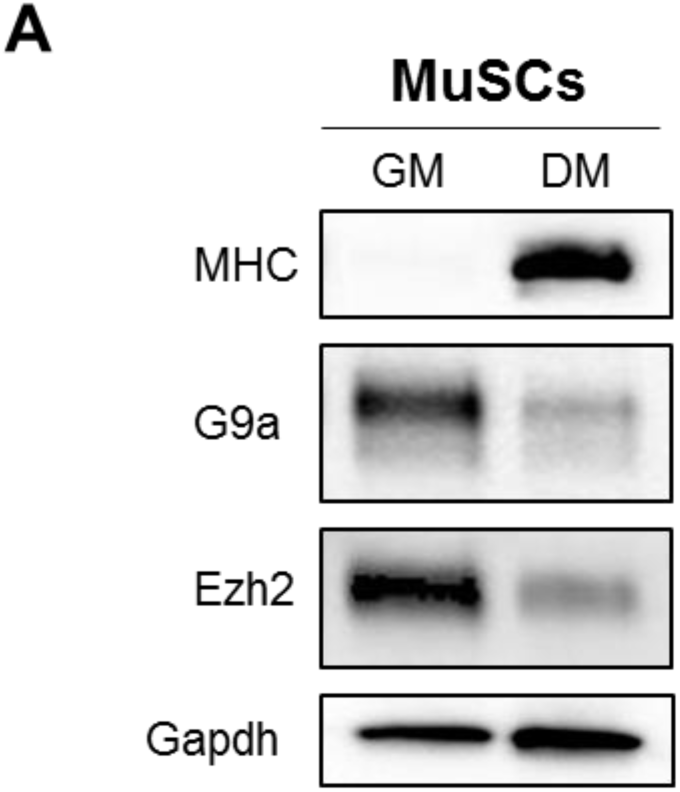
**A)** Western blot analysis performed on protein extracted from wild type MuSCs grown in growth (GM) and differentiated (DM) conditions.

## REFERENCES

Abugessaisa, I., Noguchi, S., Hasegawa, A., Harshbarger, J., Kondo, A., Lizio, M., Severin, J., Carninci, P., Kawaji, H., and Kasukawa, T. (2017). FANTOM5 CAGE profiles of human and mouse reprocessed for GRCh38 and GRCm38 genome assemblies. Sci Data 4, 170107.

Anders, S., Pyl, P.T., and Huber, W. (2015). HTSeq--a Python framework to work with high-throughput sequencing data. Bioinformatics 31, 166–169.

Babicki, S., Arndt, D., Marcu, A., Liang, Y., Grant, J.R., Maciejewski, A., and Wishart, D.S. (2016). Heatmapper: web-enabled heat mapping for all. Nucleic Acids Res 44, W147–153.

Ballarino, M., Cazzella, V., D’Andrea, D., Grassi, L., Bisceglie, L., Cipriano, A., Santini, T., Pinnaro, C., Morlando, M., Tramontano, A., et al. (2015). Novel long noncoding RNAs (lncRNAs) in myogenesis: a miR-31 overlapping lncRNA transcript controls myoblast differentiation. Mol Cell Biol 35, 728–736.

Ballarino, M., Morlando, M., Fatica, A., and Bozzoni, I. (2016). Non-coding RNAs in muscle differentiation and musculoskeletal disease. J Clin Invest 126, 2021–2030.

Batista, P.J., and Chang, H.Y. (2013). Long noncoding RNAs: cellular address codes in development and disease. Cell 152, 1298–1307.

Bolger, A.M., Lohse, M., and Usadel, B. (2014). Trimmomatic: a flexible trimmer for Illumina sequence data. Bioinformatics 30, 2114–2120.

Brack, A.S., Conboy, I.M., Conboy, M.J., Shen, J., and Rando, T.A. (2008). A temporal switch from notch to Wnt signaling in muscle stem cells is necessary for normal adult myogenesis. Cell Stem Cell 2, 50–59.

Caretti, G., Di Padova, M., Micales, B., Lyons, G.E., and Sartorelli, V. (2004). The Polycomb Ezh2 methyltransferase regulates muscle gene expression and skeletal muscle differentiation. Genes Dev 18, 2627–2638.

Carlevaro, G., Lantos, A.B., Canepa, G.E., de Los Milagros Camara, M., Somoza, M., Buscaglia, C.A., Campetella, O., and Mucci, J. (2019). Metabolic Labeling of Surface Neo-sialylglyconjugates Catalyzed by Trypanosoma cruzi trans-Sialidase. Methods Mol Biol 1955, 135–146.

Carninci, P., Kasukawa, T., Katayama, S., Gough, J., Frith, M.C., Maeda, N., Oyama, R., Ravasi, T., Lenhard, B., Wells, C., et al. (2005). The transcriptional landscape of the mammalian genome. Science 309, 1559–1563.

Chen, J., Tu, X., Esen, E., Joeng, K.S., Lin, C., Arbeit, J.M., Ruegg, M.A., Hall, M.N., Ma, L., and Long, F. (2014). WNT7B promotes bone formation in part through mTORC1. PLoS Genet 10, e1004145.

Cipriano, A., and Ballarino, M. (2018). The Ever-Evolving Concept of the Gene: The Use of RNA/Protein Experimental Techniques to Understand Genome Functions. Front Mol Biosci 5, 20.

Dobin, A., Davis, C.A., Schlesinger, F., Drenkow, J., Zaleski, C., Jha, S., Batut, P., Chaisson, M., and Gingeras, T.R. (2013). STAR: ultrafast universal RNA-seq aligner. Bioinformatics 29, 15–21.

Eden, E., Navon, R., Steinfeld, I., Lipson, D., and Yakhini, Z. (2009). GOrilla: a tool for discovery and visualization of enriched GO terms in ranked gene lists. BMC Bioinformatics 10, 48.

Eliazer, S., Muncie, J.M., Christensen, J., Sun, X., D’Urso, R.S., Weaver, V.M., and Brack, A.S. (2019). Wnt4 from the Niche Controls the Mechano-Properties and Quiescent State of Muscle Stem Cells. Cell Stem Cell.

Engreitz, J.M., Haines, J.E., Perez, E.M., Munson, G., Chen, J., Kane, M., McDonel, P.E., Guttman, M., and Lander, E.S. (2016). Local regulation of gene expression by lncRNA promoters, transcription and splicing. Nature 539, 452–455.

Fatica, A., and Bozzoni, I. (2014). Long non-coding RNAs: new players in cell differentiation and development. Nat Rev Genet 15, 7–21.

Figeac, N., and Zammit, P.S. (2015). Coordinated action of Axin1 and Axin2 suppresses beta-catenin to regulate muscle stem cell function. Cell Signal 27, 1652–1665.

Guttman, M., and Rinn, J.L. (2012). Modular regulatory principles of large non-coding RNAs. Nature 482, 339–346.

Hu, P., Chu, J., Wu, Y., Sun, L., Lv, X., Zhu, Y., Li, J., Guo, Q., Gong, C., Liu, B., et al. (2015). NBAT1 suppresses breast cancer metastasis by regulating DKK1 via PRC2. Oncotarget 6, 32410–32425.

Huarte, M., Guttman, M., Feldser, D., Garber, M., Koziol, M.J., Kenzelmann-Broz, D., Khalil, A.M., Zuk, O., Amit, I., Rabani, M., et al. (2010). A large intergenic noncoding RNA induced by p53 mediates global gene repression in the p53 response. Cell 142, 409–419.

Khalil, A.M., Guttman, M., Huarte, M., Garber, M., Raj, A., Rivea Morales, D., Thomas, K., Presser, A., Bernstein, B.E., van Oudenaarden, A., et al. (2009). Many human large intergenic noncoding RNAs associate with chromatin-modifying complexes and affect gene expression. Proc Natl Acad Sci U S A 106, 11667–11672.

Kopp, F., and Mendell, J.T. (2018). Functional Classification and Experimental Dissection of Long Noncoding RNAs. Cell 172, 393–407.

Lacour, F., Vezin, E., Bentzinger, C.F., Sincennes, M.C., Giordani, L., Ferry, A., Mitchell, R., Patel, K., Rudnicki, M.A., Chaboissier, M.C., et al. (2017). R-spondin1 Controls Muscle Cell Fusion through Dual Regulation of Antagonistic Wnt Signaling Pathways. Cell Rep 18, 2320–2330.

Le Grand, F., Jones, A.E., Seale, V., Scime, A., and Rudnicki, M.A. (2009). Wnt7a activates the planar cell polarity pathway to drive the symmetric expansion of satellite stem cells. Cell Stem Cell 4, 535–547.

Legnini, I., Di Timoteo, G., Rossi, F., Morlando, M., Briganti, F., Sthandier, O., Fatica, A., Santini, T., Andronache, A., Wade, M., et al. (2017). Circ-ZNF609 Is a Circular RNA that Can Be Translated and Functions in Myogenesis. Mol Cell 66, 22–37 e29.

Ling, B.M., Bharathy, N., Chung, T.K., Kok, W.K., Li, S., Tan, Y.H., Rao, V.K., Gopinadhan, S., Sartorelli, V., Walsh, M.J., et al. (2012). Lysine methyltransferase G9a methylates the transcription factor MyoD and regulates skeletal muscle differentiation. Proc Natl Acad Sci U S A 109, 841–846.

Lu, L., Sun, K., Chen, X., Zhao, Y., Wang, L., Zhou, L., Sun, H., and Wang, H. (2013). Genome-wide survey by ChIP-seq reveals YY1 regulation of lincRNAs in skeletal myogenesis. EMBO J 32, 2575–2588.

Maamar, H., Cabili, M.N., Rinn, J., and Raj, A. (2013). linc-HOXA1 is a noncoding RNA that represses Hoxa1 transcription in cis. Genes Dev 27, 1260–1271.

McHugh, C.A., Chen, C.K., Chow, A., Surka, C.F., Tran, C., McDonel, P., Pandya-Jones, A., Blanco, M., Burghard, C., Moradian, A., et al. (2015). The Xist lncRNA interacts directly with SHARP to silence transcription through HDAC3. Nature 521, 232–236.

Mozzetta, C. (2016). Isolation and culture of muscle stem cells, Vol 1480.

Mozzetta, C., Boyarchuk, E., Pontis, J., and Ait-Si-Ali, S. (2015). Sound of silence: the properties and functions of repressive Lys methyltransferases. Nat Rev Mol Cell Biol 16, 499–513.

Nagano, T., Mitchell, J.A., Sanz, L.A., Pauler, F.M., Ferguson-Smith, A.C., Feil, R., and Fraser, P. (2008). The Air noncoding RNA epigenetically silences transcription by targeting G9a to chromatin. Science 322, 1717–1720.

Noguchi, S., Arakawa, T., Fukuda, S., Furuno, M., Hasegawa, A., Hori, F., Ishikawa-Kato, S., Kaida, K., Kaiho, A., Kanamori-Katayama, M., et al. (2017). FANTOM5 CAGE profiles of human and mouse samples. Sci Data 4, 170112.

Nusse, R., and Clevers, H. (2017). Wnt/beta-Catenin Signaling, Disease, and Emerging Therapeutic Modalities. Cell 169, 985–999.

Ong, M.S., Cai, W., Yuan, Y., Leong, H.C., Tan, T.Z., Mohammad, A., You, M.L., Arfuso, F., Goh, B.C., Warrier, S., et al. (2017). ’Lnc’-ing Wnt in female reproductive cancers: therapeutic potential of long non-coding RNAs in Wnt signalling. Br J Pharmacol 174, 4684–4700.

Pandey, R.R., Mondal, T., Mohammad, F., Enroth, S., Redrup, L., Komorowski, J., Nagano, T., Mancini-Dinardo, D., and Kanduri, C. (2008). Kcnq1ot1 antisense noncoding RNA mediates lineage-specific transcriptional silencing through chromatin-level regulation. Mol Cell 32, 232–246.

Pang, K.C., Frith, M.C., and Mattick, J.S. (2006). Rapid evolution of noncoding RNAs: lack of conservation does not mean lack of function. Trends Genet 22, 1–5.

Parisi, A., Lacour, F., Giordani, L., Colnot, S., Maire, P., and Le Grand, F. (2015). APC is required for muscle stem cell proliferation and skeletal muscle tissue repair. J Cell Biol 210, 717–726.

Polesskaya, A., Seale, P., and Rudnicki, M.A. (2003). Wnt signaling induces the myogenic specification of resident CD45+ adult stem cells during muscle regeneration. Cell 113, 841–852.

Rinn, J.L., and Chang, H.Y. (2012). Genome regulation by long noncoding RNAs. Annu Rev Biochem 81, 145–166.

Rinn, J.L., Kertesz, M., Wang, J.K., Squazzo, S.L., Xu, X., Brugmann, S.A., Goodnough, L.H., Helms, J.A., Farnham, P.J., Segal, E., et al. (2007). Functional demarcation of active and silent chromatin domains in human HOX loci by noncoding RNAs. Cell 129, 1311–1323.

Robinson, J.T., Thorvaldsdottir, H., Winckler, W., Guttman, M., Lander, E.S., Getz, G., and Mesirov, J.P. (2011). Integrative genomics viewer. Nat Biotechnol 29, 24–26.

Rossi, F., Legnini, I., Megiorni, F., Colantoni, A., Santini, T., Morlando, M., Di Timoteo, G., Dattilo, D., Dominici, C., and Bozzoni, I. (2019). Circ-ZNF609 regulates G1-S progression in rhabdomyosarcoma. Oncogene 38, 3843–3854.

Rudolf, A., Schirwis, E., Giordani, L., Parisi, A., Lepper, C., Taketo, M.M., and Le Grand, F. (2016). beta-Catenin Activation in Muscle Progenitor Cells Regulates Tissue Repair. Cell Rep 15, 1277–1290.

Shen, P., Pichler, M., Chen, M., Calin, G.A., and Ling, H. (2017). To Wnt or Lose: The Missing Non-Coding Linc in Colorectal Cancer. Int J Mol Sci 18.

Sun, Q., Hao, Q., and Prasanth, K.V. (2018). Nuclear Long Noncoding RNAs: Key Regulators of Gene Expression. Trends Genet 34, 142–157.

Ulitsky, I., and Bartel, D.P. (2013). lincRNAs: genomics, evolution, and mechanisms. Cell 154, 26–46.

von Maltzahn, J., Chang, N.C., Bentzinger, C.F., and Rudnicki, M.A. (2012). Wnt signaling in myogenesis. Trends Cell Biol 22, 602–609.

Wang, K.C., Yang, Y.W., Liu, B., Sanyal, A., Corces-Zimmerman, R., Chen, Y., Lajoie, B.R., Protacio, A., Flynn, R.A., Gupta, R.A., et al. (2011). A long noncoding RNA maintains active chromatin to coordinate homeotic gene expression. Nature 472, 120–124.

Wang, L., Zhao, Y., Bao, X., Zhu, X., Kwok, Y.K., Sun, K., Chen, X., Huang, Y., Jauch, R., Esteban, M.A., et al. (2015). LncRNA Dum interacts with Dnmts to regulate Dppa2 expression during myogenic differentiation and muscle regeneration. Cell Res 25, 335–350.

Wang, Z., Shu, W., Lu, M.M., and Morrisey, E.E. (2005). Wnt7b activates canonical signaling in epithelial and vascular smooth muscle cells through interactions with Fzd1, Fzd10, and LRP5. Mol Cell Biol 25, 5022-5030.

Zarkou, V., Galaras, A., Giakountis, A., and Hatzis, P. (2018). Crosstalk mechanisms between the WNT signaling pathway and long non-coding RNAs. Noncoding RNA Res 3, 42–53.

Zhao, J., Sun, B.K., Erwin, J.A., Song, J.J., and Lee, J.T. (2008). Polycomb proteins targeted by a short repeat RNA to the mouse X chromosome. Science 322, 750–756.

Zhao, Y., Chen, M., Lian, D., Li, Y., Li, Y., Wang, J., Deng, S., Yu, K., and Lian, Z. (2019). Non-Coding RNA Regulates the Myogenesis of Skeletal Muscle Satellite Cells, Injury Repair and Diseases. Cells 8.

Zhu, M., Liu, J., Xiao, J., Yang, L., Cai, M., Shen, H., Chen, X., Ma, Y., Hu, S., Wang, Z., et al. (2017). Lnc-mg is a long non-coding RNA that promotes myogenesis. Nat Commun 8, 14718.

